# Dynamic Self-Reinforcement of Gene Expression Determines Acquisition and Retention of Cellular Mechanical Memory

**DOI:** 10.1101/2021.06.23.449595

**Authors:** Christopher C. Price, Jairaj Mathur, Joel D. Boerckel, Amit Pathak, Vivek B. Shenoy

## Abstract

Mechanotransduction describes activation of gene expression by changes in the cell’s physical microenvironment. Recent experiments show that mechanotransduction can lead to long-term “mechanical memory”, where cells cultured on stiff substrates for sufficient time (priming phase) maintain altered phenotype after switching to soft substrates (dissipation phase), as compared to unprimed controls. The timescale of memory acquisition and retention is orders of magnitude larger than the timescale of mechanosensitive cellular signaling, and memory retention time changes continuously with priming time. We develop a model that captures these features by accounting for positive reinforcement in mechanical signaling. The sensitivity of reinforcement represents the dynamic transcriptional state of the cell composed of protein lifetimes and 3D chromatin organization. Our model provides a single framework connecting microenvironment mechanical history to cellular outcomes ranging from no memory to terminal differentiation. Predicting cellular memory of environmental changes can help engineer cellular dynamics through changes in culture environments.

## Introduction

Cellular mechanical memory describes how cells acquire and retain information about the mechanical properties of their microenvironment. These extracellular matrix (ECM) properties impact cellular structure, function, and identity (*1*–*3*), and recent experiments suggest that this linkage depends on not just the present microenvironment but the accumulated mechanical history experienced by the cell (*4*–*10*). The mechanism by which this memory is developed, maintained, and lost is not yet understood and exhibits several unusual features. First, the timescale at which the cell responds to mechanical changes through signaling (minutes to hours) is an order of magnitude faster than the timescale of memory development and dissipation (days to weeks). This implies that microenvironmental information is rapidly acquired and used by the cell, but stored and released much more slowly. Second, the persistence time of the developed mechanical memory ranges continuously from no memory all the way to permanent memory (cell differentiation), simply by varying the microenvironmental history that the cell is exposed to (**Fig. 1A**). This strong coupling between the dynamics of memory retention and the dynamics of the stimulus being remembered is not found in common physical systems such as magnetic or shape memory materials. Understanding these unique dynamical phenomena is critical to engineering cell behavior and fate through temporal control of the cell’s physical environment.

**Fig. 1.**
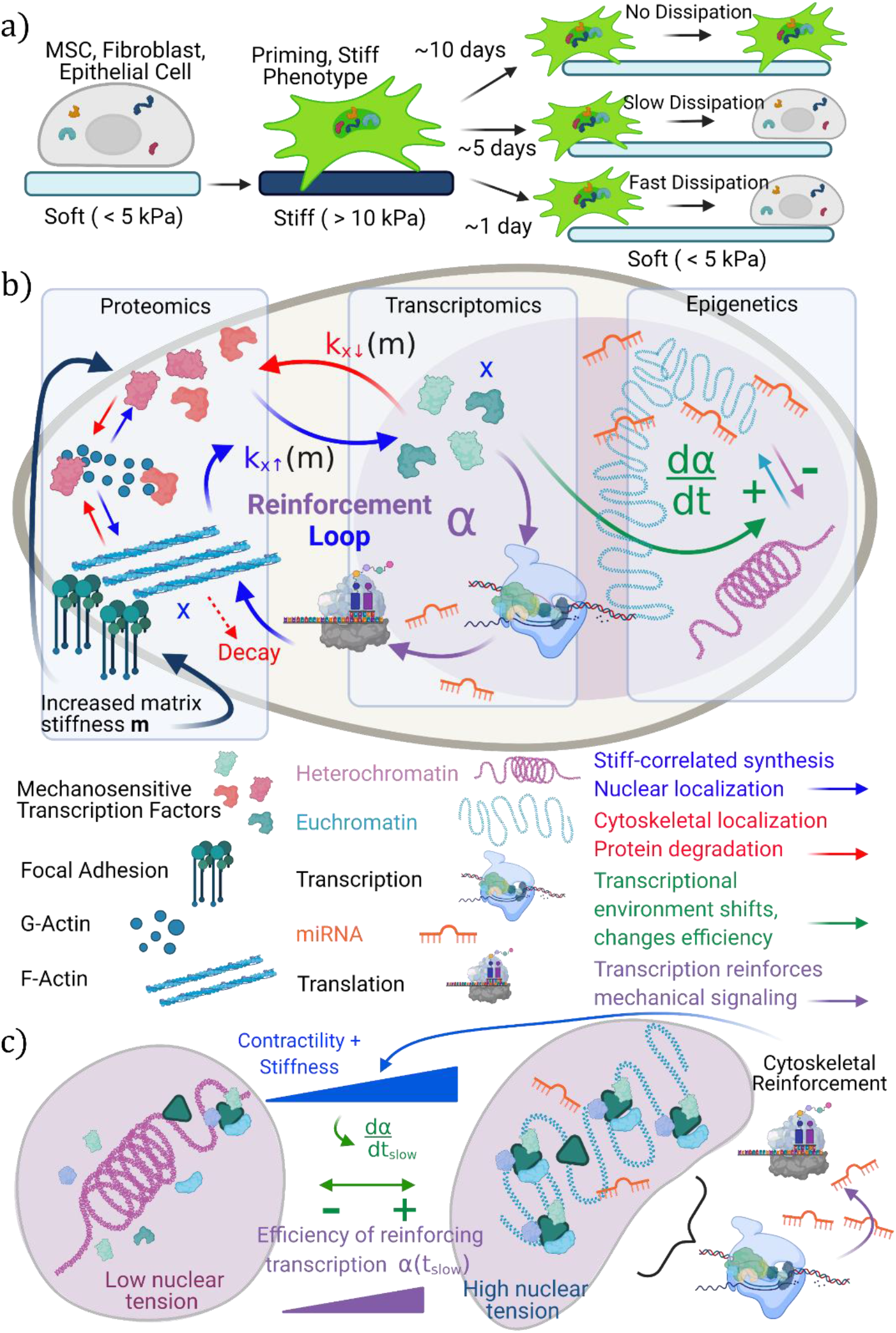
Model for dynamic mechanical memory in cells. **(A)** Experimental observations indicate that cells alter their phenotype when placed on stiff substrates (priming) in a matter of hours. The length of time that these phenotype features are retained when the cell is transitioned backed to soft substrates depends on the length of priming time on the scale of days. **(B)** Integrated cellular picture of mechanosensitive signaling and positive reinforcement enable by transcription and translation. Increased ECM stiffness leads to F actin formation, increased cellular contractility, and nuclear localization of mechanosensors (blue arrows), while soft ECM stiffness leads to decomposition of these features (red arrows). **(C)** Slow changes in the stable chromatin state in response to nuclear tension, epigenetic changes, and shifts in the post transcriptional regulation environment affect the efficiency of stiff phenotype reinforcement. High levels of reinforcement stabilize the stiff phenotype features.

Cellular adaptation to changes in the mechanical environment occurs in both the cytoskeletal and nuclear domains (*11*, *12*). On stiff substrates, examples of cytoskeletal phenotype changes include increased clustering of focal adhesions, actomyosin contractility, cell spreading area, and migration speed (*13*–*15*). On soft substrates, contractility is reduced and the mechanical properties of the cell adjust to match that of the surrounding environment by depolymerization of F-actin (*16*–*18*). In the nucleus, the population of transcriptionally active proteins changes with ECM stiffness as certain transcription factors relocate in response to mechanical signals(*19*, *20*). The chromatin structure experiences epigenetic modifications and physical deformation of the nuclear envelope from contractile forces, leading to alterations in gene expression (*21*, *22*). The dynamic nature of mechanical memory development and depletion indicates that information about microenvironmental mechanics is continuously consumed by the cell, allowing stem cell differentiation to proceed from different time series of mechanical microenvironments (*1*, *4*, *6*, *23*).

A hallmark of mathematical models of memory is bistability, which is a property of a system to have more than one steady state, and this concept forms the basis for Waddington’s famous landscape of cell differentiation. Bistability alone does not contain any information about dynamics of memory development or retention, only that it can occur (*24*, *25*). Several mechanistic models have been put forward to explain the relationship between mechanics and cell differentiation (*5*, *26*–*28*), but these models do not simultaneously capture 1) the timescale disparity between mechanical signaling / cell adaptation and memory development and 2) the continuous range of memory outcomes. More generally, regulatory gene network models with different topologies can give rise to memory using network motifs such as positive and negative reinforcement (*29*–*34*). However, explicit molecular network models for mechanotransduction are difficult to develop because there is not enough data available to determine the many model parameters or assert which components of the regulatory network are rate-limiting. This leads to rigid models which are difficult to interpret and cannot generalize across variations in priming time and priming stiffness, limiting their predictive power.

In this work, we propose a model to describe the dynamics of mechanotransductive memory acquisition and persistence. The model starts from a general molecular framework, incorporating both fast and slow mechanosensitive pathways. We simplify this model to two ordinary differential equations, representing cytoskeletal and nuclear dynamics, respectively. First, we show that simple positive reinforcement between signaling and transcription is sufficient for mechanical memory acquisition. Second, we show that dynamic coupling between the cellular phenotype and the sensitivity of this reinforcement leads to a continuous range of memory persistence time. Biologically, the sensitivity of positive reinforcement corresponds to the *epigenetic state* and *transcriptional environment* of the cell, which govern the steady-state balance between *synthesis* and *degradation* of proteins correlated with either a stiff-ECM or soft-ECM phenotype. The rate at which signaling induces changes in the positive reinforcement sensitivity (*transcriptional environment*) determines memory by shifting the phenotype *(protein composition)* from requiring external mechanical signal to a self-sustaining state. Simulating priming programs that match experimentally tested configurations, we observe emergent cases of no memory, temporary memory, and quasi-permanent memory (differentiation) by varying only the priming time and keeping other model parameters fixed. In designing future experiments or therapeutics, this simple but robust framework could help decouple the importance of positive reinforcement of mechanosensitive gene expression and their sensitivity to mechanical cues, thus optimizing the role of mechanical memory in optimizing biological outcomes.

## Results

### Model for Dynamic Mechanosensitivity in the Cytoskeleton and the Nucleus

We begin our model of mechanotransduction and mechanosensitive gene expression by introducing a matrix variable 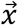 whose elements *x*_*i*=1..*n*_ represent functionally active concentrations of stiff-activated proteins and transcription factors. For a cytoskeletal protein, *x*_*i*_ corresponds to the steady-state concentration which emerges from synthesis and degradation. Examples of stiff-correlated cytoskeletal proteins include F-actin (or α-SMA), vinculin, and integrins. For transcription factors, *x*_*i*_ refers to a transcriptionally eligible concentration, which includes the steady-state level of nuclear localization. Examples of transcription factors with well-known stiff-correlated nuclear localization include YAP (*35*, *36*), MKL-1 (*20*, *37*), and RUNX2 (*38*, *39*); nuclear localization is necessary for transcription factor activity due to the possibility of co-activation requirements. We also include epigenetically modifying enzymes such as HDAC and HAT as elements of 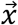 which influence chromatin organization and demonstrate mechanosensitive activity patterns (*8*). Considering all these components, the net influence of ECM stiffness on the cell quickly becomes a complex multivariate problem, and there is limited data available to inform all the cross-activity of each stiff-correlated transcription factor and protein. Therefore, we adopt a simpler approach and average the components of 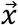 down to a single variable *x* which tracks the effective mechanoactivation over all mechanosensitive processes. However, if the matrix of interactions between the variables *x*_*i*_ is known, the present approach can be generalized as shown in **SI Section I**. The linear dynamics of *x* can be written as:

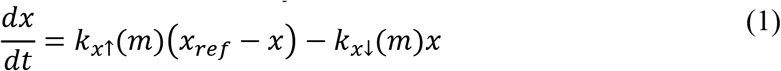

where *m* is the matrix stiffness, *k*_*x*↑_(*m*) gives the mechanosensitive rate of cytoskeletal protein synthesis and/or transcription factor nuclear import, *k*_*x*↓_(*m*) gives the rate of the reverse processes (degradation and nuclear export), and *x*_*ref*_ is a reference level of mechanoactivation at a characteristic stiffness *m*_0_. Processes described by *k*_*x*↑_(*m*) are shown with blue arrows in **Fig. 1B**, while processes described by *k*_*x*↓_(*m*) are shown with red arrows. We choose *k*_*x*↑_(*m*) to be a monotonically increasing but saturating function of stiffness, 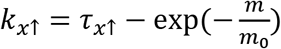, to capture the mechanosensitivity of stiff activation, and for simplicity we choose the degradation and export rate *k*_*x*↓_(*m*) to be a constant τ_*x*↓_ over stiffness (*36*) (**Fig. S1**). This is motivated by experimental evidence that nuclear import of transcription factors is more mechanosensitive than nuclear export (*36*) and that cellular response saturates at very high stiffness (*40*). While specific functional choices are arbitrary, the results we present are general to different functional forms which maintain positive correlation of *k*_*x*↑_ with stiffness. A systems circuit of our model is included in **Fig. S2** for further reference.

### Transcription Creates Positive Reinforcement Loop for Mechanical Signaling

Next, we consider that the transcriptional activity of the many components of 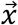 creates a positive reinforcement loop by enhancing adaptations to increased stiffness of the ECM. For example, YAP and MKL-1 activate transcription of genes which lead to increased stability of focal adhesions, F-actin, and contractility through Rho-Rock pathways and support of G-actin polymerization (*41*–*43*). This stabilization releases additional bound cytoplasmic transcription factors to translocate to the nucleus, further increasing *x*. The transcriptional positive reinforcement is depicted in **Fig. 1B** by the purple arrows; we incorporate this positive reinforcement mechanism into equation (1) by adding a nonlinear Hill relation:

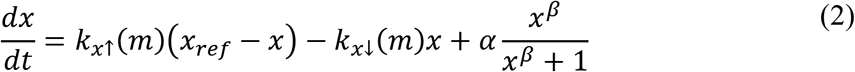

Here *α* is the sensitivity of the positive reinforcement and *β* determines the sharpness of the Hill function, which transitions from a low value to a high value like a smoothed step function. Positive reinforcement loops in cells have been extensively modeled using Hill relations and are a known source of bistability in dynamical systems (*44*–*46*). Bistability indicates at least two steady-state solutions to a dynamical system and underpins hysteresis and memory in many physical systems. Biologically, the sensitivity parameter *α* contains all the information about the efficiency of the mechanosensitive self-reinforcement, which directly corresponds to the transcription landscape. We consider this parameter as a function *α*({*y*_*i*_}, *z*), where each *y*_*i*_ represents a protein, mRNA, or non-coding miRNA involved in regulating the transcription-translation pipeline, and *z* is a label for the fraction of euchromatin relative to heterochromatin in the nucleus. Implicitly, a subset of *y*_*i*_ and *z* depend on mechanosensitive actors *x*_*i*_, which in turn depend on *m*. **Fig. 1C** illustrates how changing *α* reflects changes in both 3D chromatin architecture and post-transcriptional regulation, altering the efficiency of mechanosensitive transcription even in the limit of excess transcription factors. In the heterochromatic state, fewer chromatin sites are available for transcription. In the more active euchromatic state, a complex and modifiable regulatory environment (including miRNAs) exists in between the chromatin and downstream protein expression. Acknowledging that many transcriptional machinery and regulatory components co-depend on each other to function, we can expand *α* as a sum of terms which consider all possible contributions from coupled interactions at increasing levels of complexity.

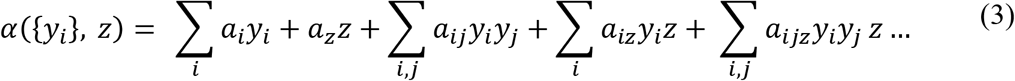

The coefficients *a*_*ij*_ act as weights which scale the relative importance of each transcriptional and regulatory component to the total state variable of the system, *α*. These weights are analogous to activity coefficients in regular solution theory, where cooperativity between different species in solution can break the linearity of mixing thermodynamics far from the dilute limit. This cooperativity arises from favorable binding interactions between solute species and long-range forces in polar media. These same features are prominent in the nucleoplasm, particularly the catalysis of transcription by formation of multi-component binding complexes (*47*, *48*).

### Fast and Slow Dynamics of Transcriptional Reinforcement Sensitivity

Since *x*_*i*_ and *m* have time dependence, we know that *α*({*y*_*i*_}, *z*) must also have a dynamic evolution which is bounded on the fast end by 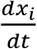 and 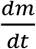 because *y_i_* shares elements of *x_i_* and *z* depends on *x*_*i*_ and *m*. Using the chain rule we can write the time derivative of *α*({*y*_*i*_}, *z*) as

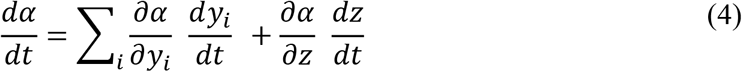

While upper-bounded by 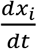 and 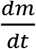, the unknown terms 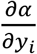 and 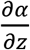 can significantly slow the overall dynamics of *α* below those of *x*_*i*_ or *m* due to complex rate-limiting or anti-cooperative relationships contained within the series expansion of *α*. Evidence of time-dependent relationships between reinforcement and transcription has been collected on individual mechanisms, including some involving mechanosensitive factors such as RUNX2 (*49*, *50*). Although we lack the data and explicit mechanistic understanding to specify *all* the contributing mechanisms to *α*, we can capture the essential nature of this time dependence by rewriting *α* as the sum of a fast changing component (on the scale of 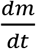 or 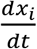) and a slow changing component which is effectively constant on the timescale of *x*_*i*_ and *m*. Complete details of the derivation are included in **SI Section I**; the result for *α*(*t*) is:

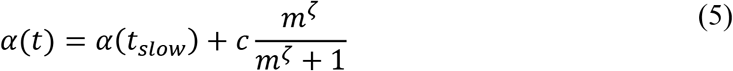

We use another Hill relation in stiffness *m* with degree *ζ* and sensitivity *c* to model the fast portion of *α*, which captures the fact that the positive reinforcement sensitivity is explicitly mechanosensitive and that stiff reinforcement requires the presence of mechanosensitive transcription factors such as YAP and MKL-1 to occur(*41*, *42*, *51*–*53*). Recent evidence indicates that the nuclear structure and chromatin conformation physically responds to environmental stiffness *via* forces transmitted through the LINC complex and not merely through chemical signals, and these direct processes are captured by this fast component of *α*(*t*) (*12*, *54*, *55*). For the remaining term *α*(*t*_*slow*_), we are free to choose a form which generally depends on *x*_*i*_ and *m* such that 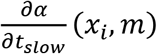 represents a weighted average of the slow, nonlinear dynamics present in equation (4).

Plugging equation (5) back into equation (2), our time-dependent equation for cellular mechanoactivation is now:

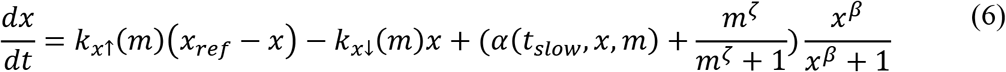

In this ODE, we established mechanosensitivity of synthesis and nuclear import of *x* (first term), mechanosensitivity of degradation and nuclear export of *x* (second term), and positive reinforcement of cellular mechanoactivation (third term) with a time-dependent sensitivity that evolves slowly with respect to changes in *x*.

### Phase Diagram of Cellular Mechanoactivation Shows Selective Bistability

We can visualize the steady-state solution space of *x* by recognizing equation (6) as the negative gradient of a “Waddington-like” energy landscape with respect to *x*, 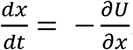. Since *α*(*t_slow_*, *x*, *m*) evolves on a much slower timescale than 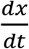, we treat *α* as a constant when finding the steady state solutions of *x*. Integrating equation (6), we arrive at

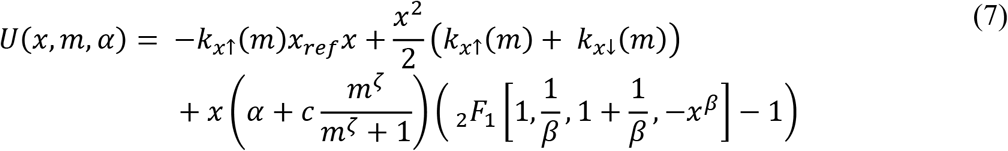

where _2_*F*_1_ is the special hypergeometric function. **Fig. 2A** gives a phase diagram of the solutions of *x* (identified as local minima in the free energy landscape) as a function of the dimensionless ECM stiffness 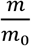 (y-axis) and the reinforcement sensitivity *α* (x-axis). The insets on the phase diagram show representative slices of the energy landscape for a point (*α*, *m*) within each region of the landscape.

**Fig. 2.**
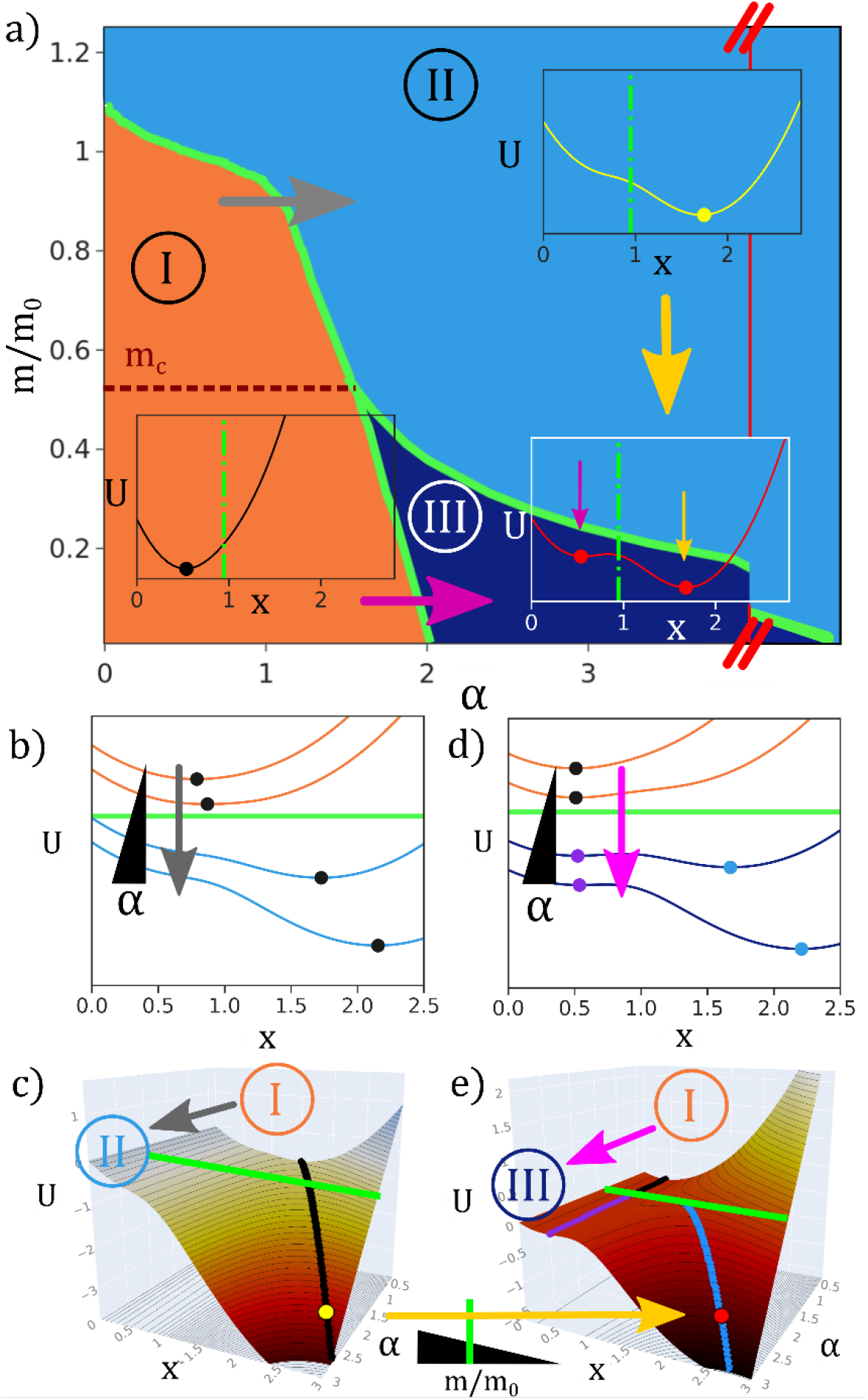
Phase diagram of the stiff-correlated phenotype. **(A)** Phase diagram of steady-state stiff phenotype expression as a function of ECM stiffness 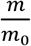 and transcriptional reinforcement sensitivity *α*. Insets demonstrate a slice of the energy surface vs. *x* for a typical point in each region, where the dots mark the energy minima and the corresponding steady state values of *x*. **(B,C)** Transitioning from region I to region II (gray arrows) by increasing *α* at constant stiffness above *m*_*c*_ leads to a significant increase in the steady-state value of *x*. Green line indicates crossing the phase boundary between regions. **(D,E)** Transitioning from region I to region III (pink arrows) at constant stiffness by increasing *α* below *m*_*c*_ traps the system in a low-*x* steady state. However, the transition from region II to region III by dropping the ECM stiffness at large *α* (gold arrows) keeps the system in a high-*x* minima.

This landscape can be divided into three regions. In orange region I (low reinforcement sensitivity and stiffness), the energy minimum and single corresponding steady state is found at small *x*. In this monostable region, there is low mechanical signal from the soft ECM, and low *α* corresponds to a small influence of the positive reinforcement process on *x*. In light blue region II, the system is still monostable, but the increased ECM stiffness induces mechanical signaling and shifts the steady-state value of *x* to a much higher value than in region I. Biologically, this corresponds to the classical pathways of mechanotransduction which occur on a timescale of hours. Compared to region I, a cell in region II exhibits greater nuclear localization of transcription factors such as YAP, RUNX2, and MKL-1, and increased focal adhesions, contractility, and areal spreading (**Fig. 1B**). Above a particular stiffness value, this shift occurs for any value of *α*. In dark blue region III (low stiffness and large reinforcement sensitivity), the system is bistable; the positive reinforcement is sufficiently strong to support a ‘stiff’ steady state at large *x* in the absence of direct mechanical signal, in addition to the expected ‘soft’ steady state at low *x*. This means that for a given value of ECM stiffness and reinforcement sensitivity in region III, the cell can exhibit either low or high mechanoactivation; this allows for hysteresis, meaning the observed state will be determined by the prior history of the ECM stiffness and the reinforcement sensitivity.

Region boundaries (green lines) in the phase diagram can be crossed by altering either the ECM stiffness or the reinforcement feedback sensitivity, inducing transitions in the steady-state mechanoactivation. Considering the soft phenotype region I as the initial condition, there are two possible transition pathways. Traversing to region II by increasing *α* above a critical stiffness (gray arrow) leads to a continuous and reversible increase in the observed value of *x* (**Fig. 2B, C**). If the mechanical signal is then removed (region II to region III, gold arrow), *x* will remain elevated as the minimum from region II smoothly transitions to the large *x* minima in region III (**Fig. 2C, E**). Traversing from region I to region III below the critical stiffness value (pink arrow, **Fig. 2D**) will not observably change *x* from the low region I value, since the region I minimum smoothly transitions to the small *x* local minimum in region III (**Fig. 2E**). Further increasing the positive reinforcement sensitivity within region III eventually leads back to region II, with a single ‘stiff’ steady state at large *x* for all values of ECM stiffness. The hysteresis loop created by the path dependence in the stiffness-reinforcement phase diagram provides a mechanism for dynamic mechanosensitive memory. A key feature of the phase diagram which corresponds to experimental observations is that increasing mechanical stiffness alone can increase *x*, allowing the cell to begin adapting to the environment on short timescales by expressing stiff-correlated proteins and localizing stiff-correlated transcription factors to the nucleus (*1*). However, these changes are fully reversible (exhibit no memory) unless the sensitivity of the positive reinforcement is sufficiently large. In the next section, we explore how evolving *α* on a slow timescale can lead to different expressions of mechanical memory depending on the time program of external mechanical stimulus.

### Nonlinear Dynamics of Positive Reinforcement Sensitivity Capture Full Range of Memory Retention Outcomes

Having shown that the trajectory of *α* can determine if memory is observed for a particular ECM mechanical history, we return to *α*(*t*_*slow*_) in equation 5 and consider an explicit form for the slow evolution of the reinforcement sensitivity. Given sufficient data on low-level biological dynamics, *α*(*t*_*slow*_) can be rigorously derived from equation (4) (**SI Section II**), but in lieu of this data, we choose the following form to maximize simplicity while capturing key phenomenological features from experiment:

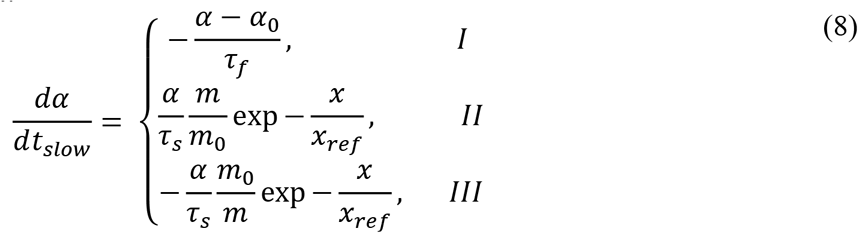

where *τ*_*f*_ and *τ*_*s*_ are time constants on the scale of hours and days, respectively, and can be directly related to 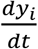 and 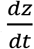 in equation (4). **Fig. 3** overlays the different piecewise components of equation (8) and their biological interpretation as reinforcement dynamics on top of the phase diagram from **Fig. 2A**. The y-axis remains the rescaled ECM stiffness *m*/*m*_0_ and the x-axis the strength of positive mechanosensitive reinforcement *α*.

**Fig. 3.**
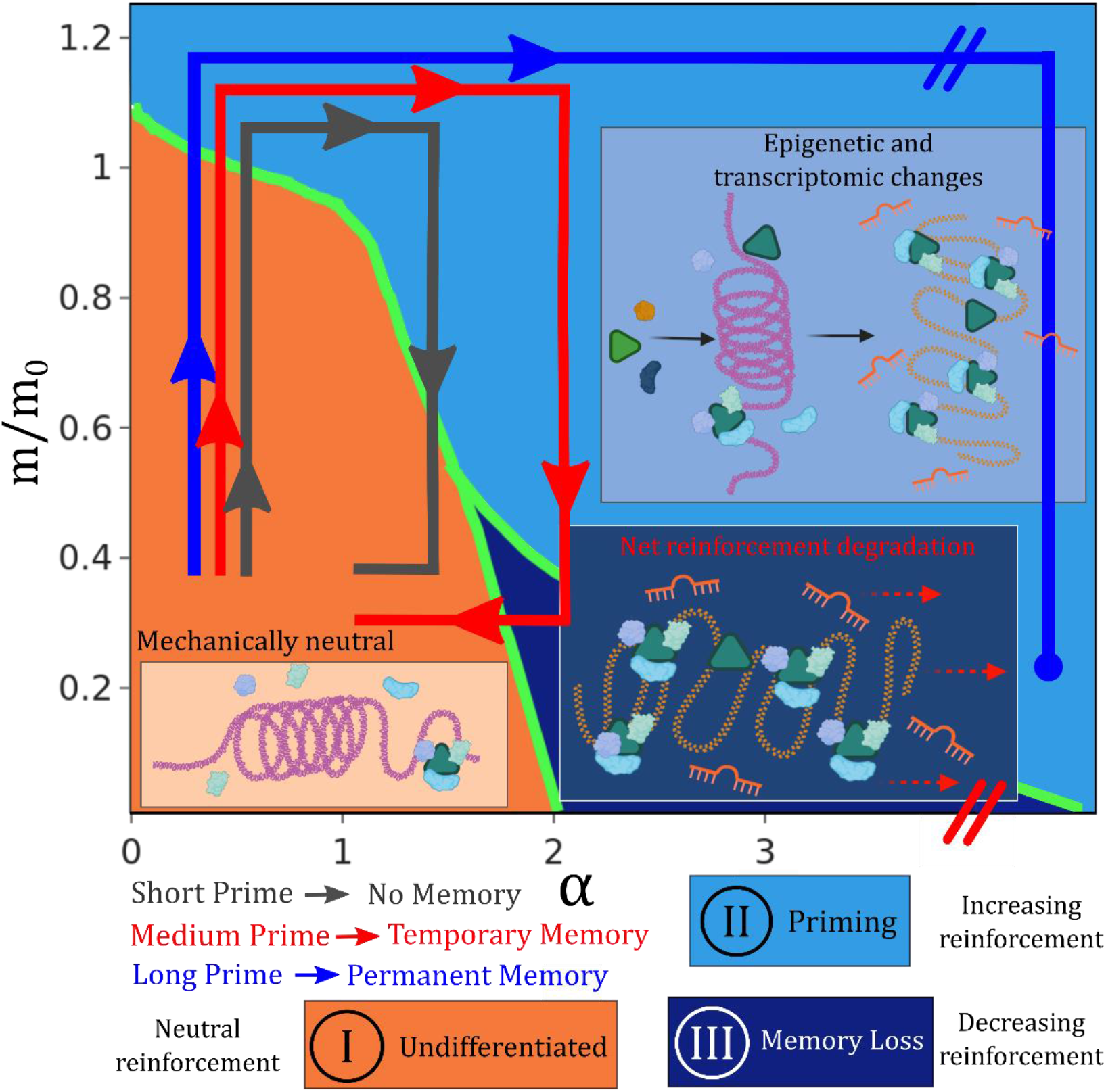
Dynamics of the transcriptional environment. In region I, the cell receives little mechanical signal and has limited positive reinforcement, so there is no driving force for the transcriptional environment to shift. In region II, signaling is sufficient to drive chromatin reorganization and changes t o the post transcriptional regulatory environment, such as miRNA synthesis. In region III, the mechanical signal is lost and there is net degradation / reversal of the stiff correlated phenotype. As self reinforcement *α* increases, less external mechanical signal is required to maintain the stiff phenotype cultivated in region II.

In region I (low stiffness and cytoskeletal reinforcement) we simply set 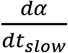 to quickly converge to a reference value *α*_0_. At low levels of mechanical signaling and without prior mechanical activation, there is no driving force to spur phenotypic change. While soft ECMs promote cell differentiation and memory, in our example we are only considering stiff-correlated genes for *x*, and there is no evidence for undifferentiated cells to develop memory which *resists* stiff priming. In **Fig. 3**, this corresponds to no change in the chromatin state or transcriptional activity over time. Memory develops at high stiffness and is lost at low stiffness unless the cell differentiates, so we choose *α* to increase in region II and decrease in region III to complete our piecewise description. By our definition, increasing *α*(*t*_*slow*_) in region II accounts for slow, nonlinear processes (shifts in the 3D chromatin and transcriptional regulation environment) which increase reinforcement of a stiff cellular phenotype (**Fig. 3**). Decreasing *α*(*t*_*slow*_) in region III models net decay of these stiff phenotype features (which can have lifetimes on the scale of days to weeks (*56*)) and reversal of the transcriptional environment in the absence of sufficient mechanical signal.

In the priming region II, multiplying by 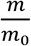 ensures that the priming time required to achieve a given level of memory decreases when increasing the priming stiffness. Including a dependence on *x* ensures that the persistence time of mechanical memory increases nonlinearly with priming time for a specified priming stiffness (*6*). Mechanistically, our definition of *x* includes mechanosensitive epigenetic modifiers such as HDAC and HAT (*8*, *57*, *58*), and while the activity of these enzymes to flip epigenetic marks occurs on shorter timescales relative to memory (*57*), chromatin structural organization and downstream effects on transcription can be much slower due to glassy dynamics of actual chromatin conformational change (*59*–*61*). This couples the slow dynamics of the reinforcement sensitivity to the steady-state value of *x*, which changes depending on the specific location within each region of the phase diagram. This coupling of reinforcement sensitivity to the signal itself is a new feature of our model which has not been studied in other models of cellular positive reinforcement loops. For simplicity and to limit free parameters, we choose the *α* degradation dynamics in region III to be the reverse of the priming dynamics. Net degradation of the reinforcement and dissipation of memory will be faster at smaller *m* and will smoothly change from the value of *α* in region II.

Each of the three arrows (grey, red, and blue) in **Fig. 3** correspond to a different hypothetical stiff priming program which leads to a different class of memory outcome. The initial conditions, priming stiffness, and model parameters (Table 1) are fixed across the three programs. The corresponding time evolution of *x* and *α* for each mechanical priming program is plotted in **Fig. 4A-C**. Between each of the three priming program results shown in **Fig. 4A-C**, only the length of time that the simulated cell is exposed to stiff substrate (10 kPa) is changed; the soft substrate is modeled at 2 kPa.

**Fig. 4.**
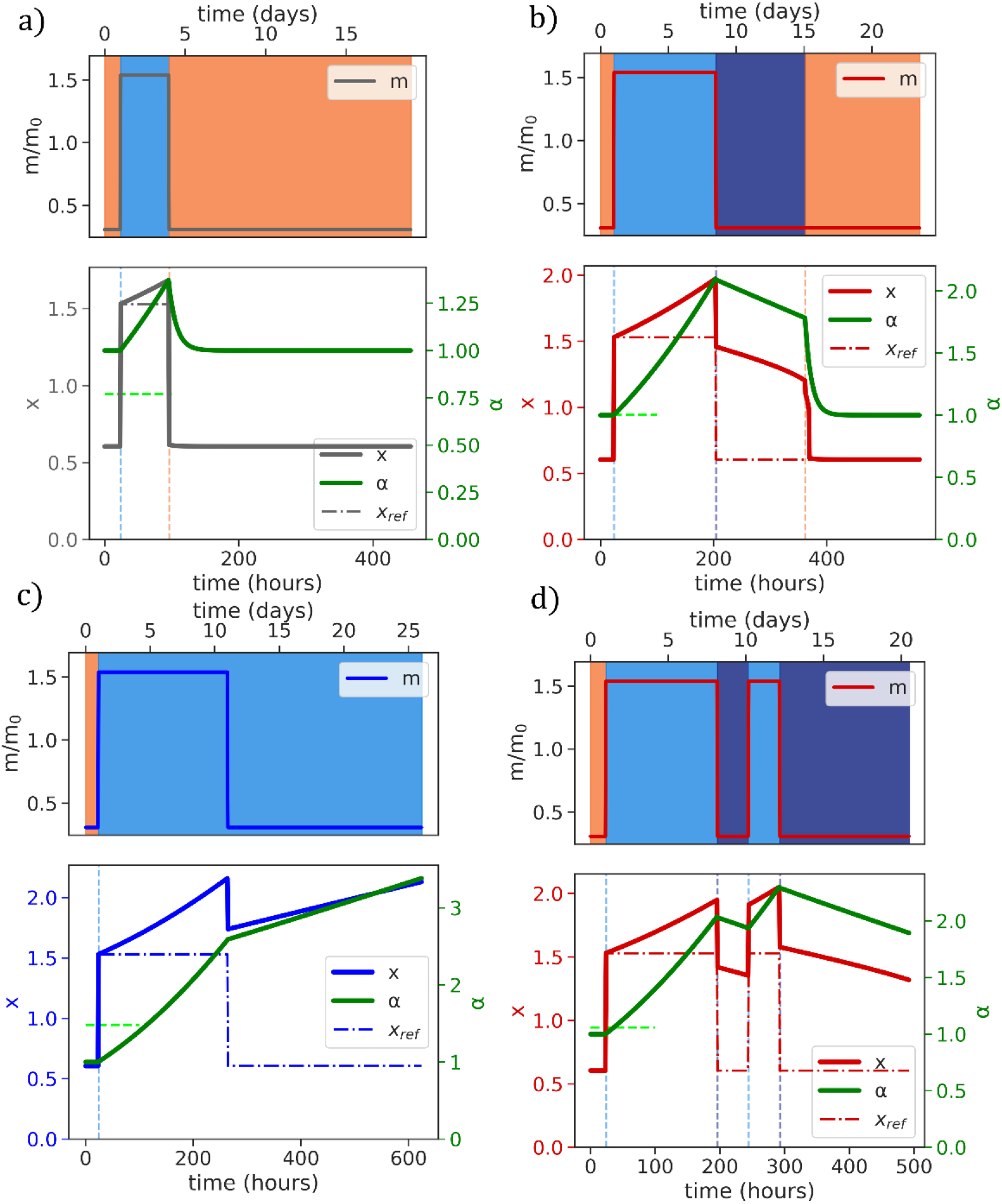
Applying different mechanical priming programs. Dot-dash lines *x*_*ref*_ indicate the value of *x* without *α* dynamics (*α* = *α*_0_). **(A)** Short priming time of a few days does not result in memory, corresponding to the gray trajectory in (D) and in **Fig. 3**. **(B)** Medium priming time results in memory on the timescale of priming, but eventually this memory decays and the system resets. **(C)** Longer priming time prevents the system from entering the memory dissipation region when the substrate stiffness is decreased, leading to permanent memory of stiff phenotype. All model parameters in (A-C) are fixed except for the length of priming time in the mechanical program (top plots). **(D)** Two-phase mechanical priming program which illustrates cumulative priming. The first priming phase is identical to **(B)**, and the total priming is equivalent to **(C)**. The second short priming pulse generates significantly more memory than the first priming pulse, yet the cell remains reversibly plastic compared to (C) since some priming is reversed between the two pulses.

**Table 1.** Parameters for simulations in Fig. 4A-D.

| Parameters (Fig. 4) | Values | Units |
| --- | --- | --- |
| Phase Diagram |  |  |
| $m_0$ | 6.5 | kPa |
| $x_{ref}$ | 2. | Arb. |
| $\beta$ | 6 | n/a |
| $\zeta$ | 35 | n/a |
| $c$ | 1. | Hours<sup>-1</sup> |
| $\tau_{x\downarrow}$ | 1.5 | Hours |
| $\tau_{x\uparrow}$ | 1.5 | Hours |
| Dynamics |  |  |
| $\tau_f$ | 12. | Hours |
| $\tau_s$ | 150. | Hours |
| $\alpha_0$ | 1. | Arb. |
| Priming |  |  |
| $m_{stiff}$ | 10 | kPa |
| $m_{soft}$ | 2 | kPa |

### Short Priming Does Not Lead to Memory Formation

The grey program does not exhibit any memory – the time that the cell is exposed to the stiff environment is short, and when the cell is returned to a soft ECM, the system returns to region I. While the phenotype quickly shifts to respond to the stiffening substrate at the beginning of the priming program (crossing the dashed green line corresponding to the boundary between regions I and II), the mechanical signal is not maintained long enough to alter the transcriptional environment to the point where it can sustain memory. The stiff phenotype is lost just as rapidly as it was gained (timescale of hours) since the dynamic trajectory returns directly to region I when the stiffness is relaxed. In the case of a stem cell, this corresponds to an insufficient mechanical signal to sustain differentiation.

### Medium Priming Leads to Temporary Memory with Variable Retention Time

The red program in **Fig. 3** exhibits temporary memory – by holding the cell in priming region II for longer than the gray program, *α* increases sufficiently such that when the cell is returned to a soft environment, it enters the bistable region III. The positive reinforcement loop traps the system in a steady state of large *x* despite the absence of persisting stiff mechanical signaling (**Fig. 4B**). The dot-dash line *x*_*ref*_ shows the phenotype expression of *x* in the absence of *α* dynamics (*α* is fixed at *α*_0_) under the same priming program. The significant deviation of *x* from *x*_*ref*_ represents the ‘phenotypic distance’ of the cell from the low reinforcement case; the length of time that this difference is maintained (while the cell is in region III) gives the length of time of observed memory. Since the dynamic evolution of *α* fundamentally changes the energy surface, the persistence time of memory is decoupled from the relaxation rate of *x*, as is observed experimentally. Depending on the specific length of priming time and priming stiffness, the model predicts a continuous range of memory persistence times from much shorter than the priming time to much longer than the priming time using the same parameter set. Over time, *α* slowly decreases (driving *x* to decrease) due to the absence of continued signaling promoting epigenetic change and natural degradation of stiff phenotype proteins, dissipating memory and eventually returning the system to region I. The model also predicts that as substrate stiffness decreases after priming, the window of reversible memory (range of *α* which corresponds to region III) grows significantly. This means that the phenotype of the cell is more likely to be reversible if the dissipative mechanical signal is stronger.

### Long Priming Leads to Permanent Memory

Finally, the blue program corresponds to permanent memory, which in the case of MSCs indicates lineage specification to a stiff phenotype (osteocyte). As the sensitivity of the positive reinforcement *α* continues to increase, it requires a stronger reversing signal (softer ECM) to enter the bistable, temporary memory regime. At a certain point (beyond the axis break in **Fig. 3**), it becomes practically impossible to sufficiently reverse the mechanical signaling and the cell will permanently exhibit a phenotype correlated with large *x* and saturated large *α*. *In vitro* experiments confirm that differentiated osteocytes exhibit sustained higher nuclear activation of YAP/TAZ and other stiff-correlated proteins, qualitatively agreeing with our picture of a phenotype which retains features of high *x* (*52*). **Fig. 4C** shows how simply increasing the priming time using the same ECM stiffnesses of the mechanical programs in **Fig. 4A** and **4B** prevents the system from leaving region II of the phase diagram after the priming phase. Physically, this means that the transcriptional and epigenetic state of the cell has absorbed enough mechanical signal during the priming phase to self-sustain the stiff phenotype once that signal is removed. Even after reducing the ECM stiffness, *α* and *x* will continue to slowly increase until they reach a saturation value which corresponds to lineage specification (**Fig. S3**). The model predicts that this transition to a ‘permanent’ phenotype is a result of the net cumulative mechanical signal absorbed by the cell; for example, consecutive short priming programs will have an additive effect due to the dynamics of *α* in regions II and III (**Fig. 4D**). In this trajectory, the initial priming period is the same as that in **Fig. 4B,** but the short second prime ends up building significantly longer memory than in **Fig. 4B** due to the accumulated ‘environmental knowledge’ which is not dissipated in the short intermediate soft period. This agrees with experimental evidence that cyclical stretching and stress stiffening of cellular substrates induces stiff differentiation (a ‘pumping’ effect) (*62*–*64*). The model also predicts that if the epigenetic / transcriptional state labeled by *α* is manipulated by a drug or other mechanism, the cell can lose its permanent mechanical memory and be ‘reprogrammed’, which corresponds physically to reversible lineage specification enabled by so-called Yamanaka factors (*65*).

### Noise in *α* Dynamics Results in Qualitatively Accurate Memory Distributions when Compared with Experiment

We have so far identified and predicted a wide range of phenomenological features of cellular mechanical memory with our simple, dynamic positive reinforcement model at the single cell level. However, biological systems are inherently noisy and experimental measurements of cellular phenotype and mechanical memory are most often taken over a population of cells. We categorize possible random fluctuations in our model into two categories – noise which affects mechanosensation and signaling (‘fast’ noise) and noise which affects the slower dynamics of reinforcement (‘slow’ noise). ‘Fast’ noise contains all the fluctuations which might cause the phenotype of a cell to not occur at the local minimum of the energy landscape on fast timescales (deviations away from steady state). This is particularly relevant in the bistable region III, where fluctuations could cause cells to jump between different local minima, corresponding to changes in phenotype and changing observations of memory. In a bistable energy landscape, a normal distribution of fluctuations away from steady state values of *x* will bias a population towards the global minimum over the local minimum since the jump rate will be higher if the energy barrier height between wells is lower (**Fig. S4**). **Fig. 5A** shows the global minimum steady state value of *x* over the stiffness-reinforcement phase diagram from **Fig. 2A** (region boundaries in green). In the majority of region III, the high-*x* minimum is lower in energy. For our purposes of stiff priming programs which enter region III from a single-minima, high-*x* state in region II, this means that fluctuations from steady state will tend to reinforce a noisy population to remain in the high-*x* state, preserving memory and having little qualitative effect on the model results.

**Fig. 5.**
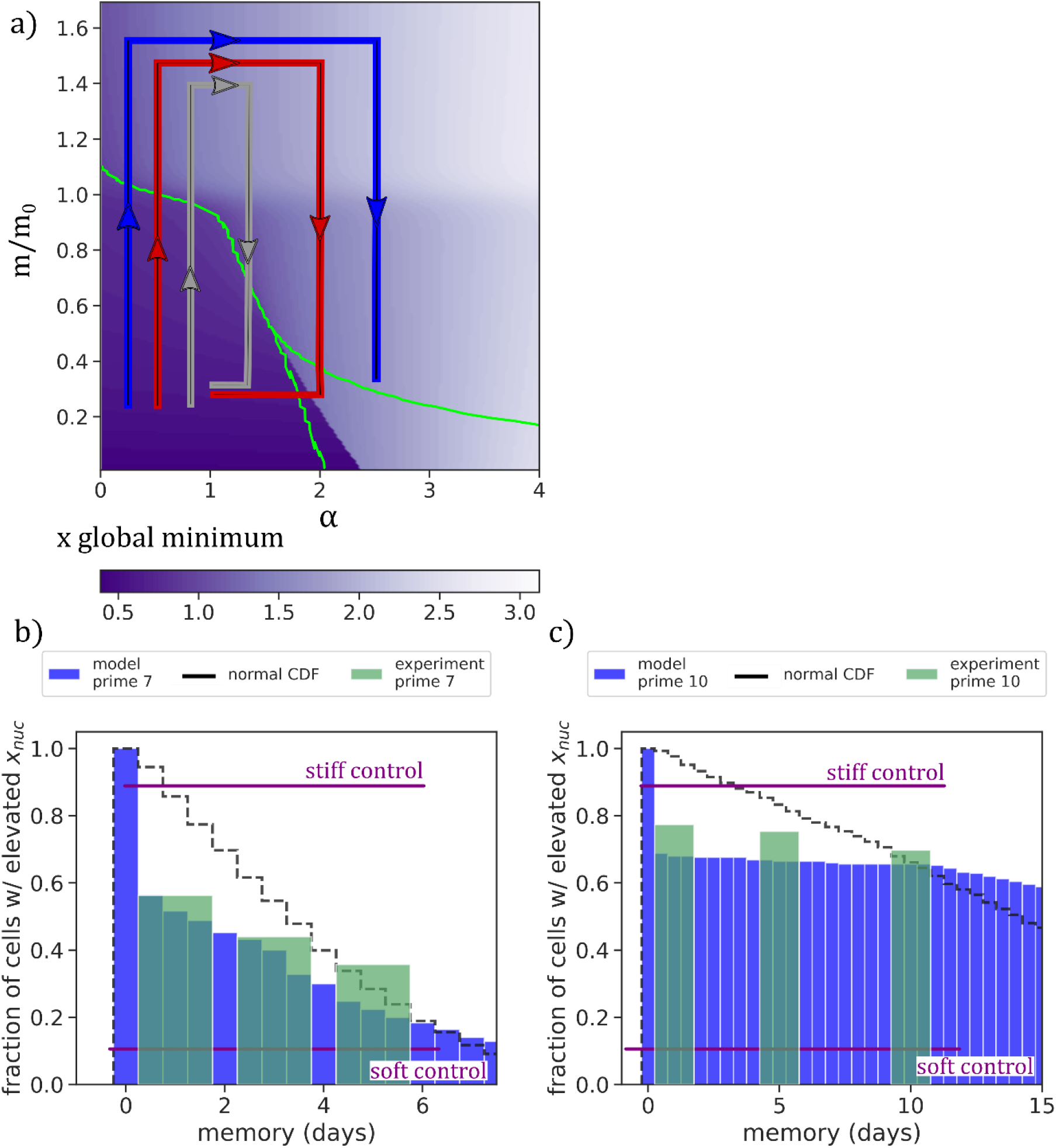
Adding noise to nonlinear *α* dynamics. **(A)** Global energy minima of *x* vs. *α* and *m* overlaid with priming programs from Fig. 3 and phase boundaries from Fig. 2. For the majority of Region III, the large x minima is also the global energy minima, indicating that noise on fast timescales is unlikely to cause well hopping and disrupt temporary memory. **(B,C)** Cumulative distribution (CDF) of memory times from simulations with slow, gaussian noise incorporated onto 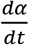 for priming of 7 days (**(B)** and 10 days **(C)**), matching experimental conditions from Yang et al. Fig. 3 (green bars and purple control lines). The black dashed line shows the CDF of a normal distribution with the same mean and standard deviation as the model distribution for reference.

‘Slow’ noise captures fluctuations in the dynamic evolution of *α*, and this is more interesting to consider due to the nonlinearity of *α*(*t*). The fact that 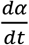 depends on the current steady state of *x* (and therefore the prior history of 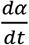) means that a normal distribution of noise in the dynamics of *α* could lead to a non-normal distribution of memory results. We investigated the impact of including noise on 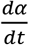 by introducing a normal distribution of noise with 0 mean, unit standard deviation, and magnitude *A*=0.01 at each time step of the simulations conducted in **Fig. 4A-C** and generating a distribution of results over N = 256 simulations. The distribution of memory times observed from the noisy simulations is shown in blue in **Fig. 5B, C**. This data contains all simulation runs including those without memory, so the difference between the first bin and second bar shows the percentage of trials (cells) which did not exhibit any mechanical memory. The thin black line gives the cumulative distribution function of a normal distribution with the same mean and standard deviation as our generated dataset. This confirms that applying normally distributed noise to the dynamic evolution of *α* results in a non-normal distribution of observed memory persistence times due to the non-linearity of the *α* dynamics.

Experimental data on persistence time of YAP and RUNX2 nuclear localization as a function of priming time on stiff substrates (10 kPa) taken from (*6*) is overlaid on **Fig. 5B, C**. We averaged their results from YAP and RUNX2 to get a general sense of how the mechanoactivated cell population changes over time (green bars) after the substrate is switched from stiff to soft (2 kPa). The purple control lines indicate the experimental baseline of mechanoactivation in nuclear localized YAP and RUNX2 without any substrate switching. With added noise, our model captures the qualitative changes in the phenotype distribution over time as priming time is changed, with longer-primed cells being more resistant to return to the soft control phenotype. As in **Fig. 4**, all parameters aside from priming time are held constant between the **Fig. 5B** and **Fig. 5C** to best replicate the experimental conditions (Table 2). In both the experimental data and the model, 10 days of priming leads to significantly higher retention of the stiff phenotype in the cell population than 7 days of priming.

**Table 2.** Parameters for simulations in Fig. 5B-C.

| Parameters (Fig. 5) | Values | Units |
| --- | --- | --- |
| Phase Diagram |  |  |
| $m_0$ | 6.1 | kPa |
| $x_{ref}$ | 1.2 | Arb. |
| $\beta$ | 4.9 | n/a |
| $\zeta$ | 35 | n/a |
| $c$ | 1 | Hours<sup>-1</sup> |
| $\tau_{x\downarrow}$ | 1.1 | Hours |
| $\tau_{x\uparrow}$ | 1.5 | Hours |
| Dynamics |  |  |
| $\tau_f$ | 12. | Hours |
| $\tau_s$ | 160. | Hours |
| $\alpha_0$ | 1. | Arb. |
| Noise |  |  |
| $\sigma$ | 0.7 | Arb. |
| $A$ | 0.01 | Arb. |
| Priming |  |  |
| $m_{stiff}$ | 10 | kPa |
| $m_{soft}$ | 2 | kPa |

To isolate the effect of the nonlinear coupling between mechanical signaling (*x*) and transcriptional environment dynamics *α*(*t*) on the population statistics, we attempted the same noisy simulations using a linear form for 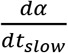 without *x* or *m* dependence (**Fig. S5**). This emphasizes that the nonlinear coupling between mechanical signaling and the dynamic evolution of the transcriptional environment is a fundamental conceptual ingredient which can explain both the disparate timescales of cellular adaptation and memory and captures non-normal population statistics. The linear noise simulations can still result in zero, temporary, or permanent memory. However, the population statistics reflect the normal distribution of the noise applied, as seen by the agreement between the red model results and the black normal distribution CDF. While the available experimental data is limited, the same set of parameters using linear dynamics cannot qualitatively capture the experimental population distribution change with priming time nearly as well as the nonlinear dynamics, despite the same number of free parameters. The selection procedure for choosing the free parameters is discussed in the Methods section.

### Model Feature Comparison with General Experimental Observations

We selected the data from the Yang et al. study on mesenchymal stem cells for direct comparison with our model since this is one of the few experimental studies to explicitly track components of cellular mechanotransduction as a function of mechanical priming time. While drawing quantitative comparisons across different experimental studies is difficult due to confounding variables such as cell lineage and growth media, we highlight several features of our model which appear in other studies (results summarized in **Table S1**). In our model, increasing ECM stiffness enough will always lead to cellular expression of a stiff phenotype on the scale of τ*x*↓ and τ*x*↑ irrespective of memory formation; our chosen values for these parameters are based on the adaptation time observed experimentally of ~1 hour (*17*). The characteristic stiffness value *m*_0_which we use in **Figure 4** and **Figure 5** is consistent with the priming and memory stiffnesses used in other experiments in **Table S1**. Short priming of ~1 day does not lead to appreciable memory in both our model and experiments (*7*), and temporary memory retention time is generally greater than or equal to the priming time across different experiments. In our phase diagram, reduction of *α* from region II to region III or region I erases permanent memory; experimentally, knockdown of miR-21 (a component of *α*(*t*_*slow*_)) also erased permanent memory even after long priming (*5*). Temporary memory development correlated with RUNX2 nuclear localization using stiff and soft substrates of 8 and 0.5 kPa after 7 days of priming was recently observed by Watson *et al.* in epithelial cells (*66*); these values are similar to the data from Yang *et al.*, indicating that similar parameters in our model are translatable to a different cell type. Finally, in our model the reinforcement strength and acquired memory is cumulative; this agrees qualitatively with experiments which have investigated dynamic cyclical stretching as a way to observe mechanical memory (*63*, *64*). **Fig. 6A-C** gives a schematic overview of the progression from external mechanical signal to self-sustaining mechanical memory by way of increased transcriptional reinforcement spurred by mechanotransduction.

**Fig. 6.**
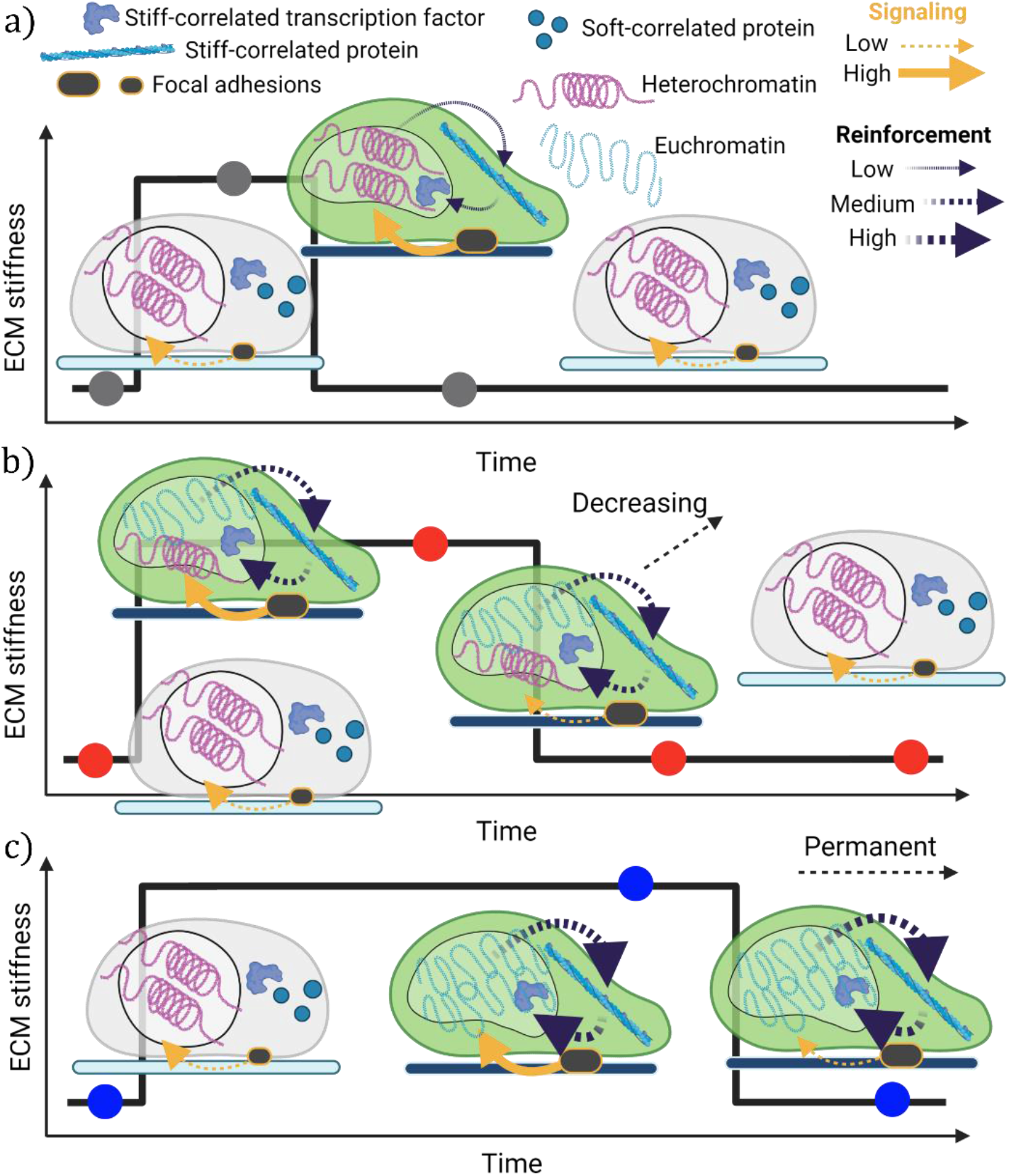
Summary of dynamic mechanical memory. **(A)** At short priming times, mechanical signaling leads to cellular adaptation but does not persist for sufficient time to increase reinforcement, leading to no memory. subsequent reinforcement is low, preventing observation of memory. **(B)** At intermediate priming times, reinforcement increases with persisting mechanical signal. The transcriptional environment shifts enough to build temporary memory, but this reinforcement will slowly decay to erase memory once the mechanical signal is removed. **(C)** At long priming times, reinforcement strength continues to grow with input mechanical signal and an adapting transcriptional environment. Reinforcement becomes strong enough to sustain without any mechanical signal, and the new phenotype persists if the substrate is changed (permanent memory).

### Simple Generalization for Analogous Soft-ECM Correlated Mechanical Memory

In this work, we focused on stiff-priming and stiff-correlated mechanical memory since these conditions are the most widely studied due to applicability in stem cell therapies for fibrosis and osteogenesis. However, cells can also develop analogous soft-correlated mechanical memory which can eventually lead to soft tissue generation such as neurogenesis with sufficient priming (*1*). Our model is instantly generalizable to this case by reconstructing *x* as an averaged quantity of soft-activated phenotype components 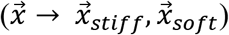 and inverting the scaled stiffness from 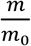 to 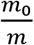 (**Fig. 6A-C**). The phase diagram for soft-correlated memory and phenotypic activation is shown in **Fig. S6** and retains the three distinct regions which allow for no memory, temporary memory, and permanent memory depending on priming time. Recent experiments which primed adipose stem cells on 1 kPa substrates for two weeks found that temporary soft memory develops with similar persistence times (between one and two weeks) to stiff memory (*10*). In contrast with stiff priming, nuclear YAP localization was not found to be a marker of soft-priming. This observation agrees well with our definition of the mechanically correlated phenotype fingerprint vector 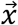; nuclear YAP is an element of the stiff-correlated 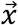 but not the soft-correlated 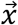. Using this simple, modular model framework, more complex models can be assembled which simultaneously consider soft and stiff memory and downstream consequences for differentiation.

## Discussion

The acquisition and maintenance of mechanical memory is a general phenomenon across different cell types and culture environments (*4*–*10*, *67*). Balestrini et. al. cultured lung fibroblasts for two weeks on stiff (100 kPa, priming phase) substrates and found that they continued to express elevated fibrotic activity after being transferred to soft substrates (5 kPa, dissipative phase) for at least two weeks. A follow-up study by Li et al. under similar conditions identified miR-21 as a necessary molecule for long-time memory maintenance, indicating the role of transcriptional efficiency in memory regulation (*5*). Some miRNAs can have half-lives on the scale of multiple days, which motivated our formulation of *α*(*t*_*slow*_) to conceptually include these non-coding RNA molecules. More recent experiments have focused on detailed changes of chromatin organization within the nucleus, confirming that epigenetic changes occur in response to mechanical signaling (*8*) and highlighting the role of the LINC complex as a direct, physical mechanosensory (*12*, *54*, *68*). Additionally, we have recently shown that *epithelial* cell sheets primed on a stiff matrix for 3 days also store mechanical memory through nuclear YAP localization, which continues to enhance cell migration through enhanced pMLC expression and focal adhesion formation on soft matrix for 2-3 days (*7*).

In developing our model, we sought to synthesize and distill the phenomenological observations from these experiments and related studies covering the impact of mechanics on lineage specification, which has not been accomplished by existing models to our knowledge. Li et. al. proposed a reservoir model along with their identification of miR-21 as a memory regulator, where priming leads to production of memory keepers which slowly dissipate after priming halts. This model alone does not explain the timescale disparity between mechanical adaptation and development of memory. Mousavi et al. and Peng et al. proposed two different mechanically activated differentiation models based on population dynamics and gene regulatory networks, but these models do not capture the variable rates of memory dissipation observed in experiments. These models rely on ~20 and ~40 free kinetic parameters, respectively, yet do not account for key qualitative features of the memory phenomena. Our model uses 8 unique free parameters, which sacrifices resolution on specific biological mechanisms but allows us to identify that a simple nonlinear coupling between signaling and transcriptional evolution is sufficient to capture the phenomenological features of cellular plasticity.

The continuous range of cellular plasticity persistence time from zero (no memory, **Fig. 6A**) to permanent (lineage specification, **Fig. 6C**) is unique when compared to other physical memory systems, which often either exhibit permanent memory or no memory. Although early studies of lineage specification viewed this process as uni-directional (such as the traditional Waddington landscape), the targeted reversibility of plasticity under the right conditions is also a unique physical feature. The traditional Waddington landscape identifies specific branch points which split cells into separate wells representing stable phenotypes (*24*). Our model generalizes this picture by showing that both the Waddington landscape surface and the rate at which the cell progresses down each well can be altered by external stimuli such as stiffness. This ‘graduated reversibility’ may function biologically to make the cell more resilient to local short-term fluctuations in environment, while still allowing for long term, correlated population shifts in response to persistent environmental cues.

Predicting the memory response of cells to their mechanical environment has significant implications for designing cell-based therapy and studying other cellular mechanisms *in vitro*. Based on our model, we predict that small changes in priming stiffness or priming time can have large consequences on the retention time of developed phenotypes due to the nonlinearity of slow-evolving components. Our phase diagram indicates that recovery of stem-like, soft phenotypes can be enhanced after priming by reducing the stiffness of the recovery substrate, extending the range of region III which allows for memory dissipation. However, beyond a certain point, mechanical signal alone will not lead to phenotype reversal due to formation of permanent memory. Measuring the extent of priming may require nuclear information and not just data on signal activity, since the timescale of signaling is independent of the timescale of memory development. External methods to change *α*, such as Yamanaka factors or changes in growth media, can overwrite the natural permanent persistence of the stiff phenotype in these situations. In future work, we anticipate that this model framework for mechanical memory can be extended to include a chemical axis, which can be used to consider more general cases of cell differentiation and coupling between chemical and mechanical contributions to memory acquisition and retention.

## Limitations

In our model, we made two key assumptions: 1) Positive feedback loops exist in mechanosensing pathways, and 2) Shifts in the transcriptional environment which affect these feedback loops depend on signaling but occur on slow timescales. Quantitative predictions of cell responses will require more experimental data to validate more complex and precise models. Based on our results in this work, determining the rate of change of the transcriptional environment (*α*(*t*)) as a function of priming stiffness and priming time is the most important unknown quantity. This is difficult to assess from epigenetic modifiers alone since chromatin reorganization occurs on a longer timescale than epigenetic enzyme activity. Hi-C experiments during both priming and memory dissipation would provide information on the rate of change of the chromatin conformation. While this would not completely specify the transcriptional environment, this information would be key to understand which steps are rate-limiting in the evolution of mechanically activated cellular plasticity. The further that *α*(*t*) can be specified with mechanistic information from the nucleus, the greater predictive accuracy on the dynamics of memory can be.

## Materials and Methods

The model was implemented using a standard ODE solver (fsolve) in the open source SciPy package (Python). In **Fig. 5**, parameter selection was done using a design-of-experiments approach. A latin hypercube sampling method was used to generate combinations of the parameters *α*_0_, *m*_0_, *x*_*ref*_, τ_*s*_, τ_*x*↓_, τ_*x*↑_, *n*, *σ*, and *A*. Each parameter combination was run for priming times of 3, 7, and 10 days, with 250 noisy trials run for each priming time. Parameter combinations were scored against the experimental data from Yang *et* al. using a Kolmogorov-Smirnov test, a least-squares test, and manual inspection. The data shown in **Fig. 5** (nonlinear dynamics) and **Fig. S6** (linear dynamics) represent the distributions with the best-fit from the sampling conducted in both metrics. We note that these do not represent best-fits to the data but were sufficient to show qualitative agreement and differentiate the two different dynamics approaches.

## Acknowledgements

This work was supported by National Cancer Institute award R01CA232256 (VBS), National Institute of Biomedical Imaging and Bioengineering awards R01EB017753 (VBS) and R01EB030876 (VBS), NSF Center for Engineering Mechanobiology grant CMMI-154857 (JB, AP & VBS), NSF grants MRSEC/DMR-1720530 (VBS) and DMS-1953572 (VBS), NIH/NIGMS award R35GM128764 (AP), and NIAMS grant 5R01AR073809 (JB).

