## Supporting Information for "Dynamic Self-Reinforcement of Gene Expression Determines Acquisition and Retention of Cellular Mechanical Memory"

**This PDF file includes:**

Supplementary Text
Figs. S1 to S6
Table S1

**Supplementary Text**

Section I: Generalized Model for Dynamic Self-Reinforcing Mechanosensitivity

Consider a vector variable $\vec{x}$ of dimension $n$ where each element $x_i$ represents the nuclear concentration of a stiff mechanosensitive transcription factor or the cytoskeletal concentration of a stiff phenotype protein. $\vec{x}$ is a fingerprint state vector for the mechanical phenotype of the cell. For each element of $\vec{x}$, we can write a linear steady-state rate equation

$$\dot{x}_i = k_{\uparrow i}(m)\left(x_i^{ref} - x_i\right) - k_{\downarrow i}(m)x_i + \sum_j c_{ij}(m)x_i x_j \quad \text{(s1)}$$

where $\dot{x}_i = \frac{dx_i}{dt}$ , $m$ is the ECM stiffness, $k_{\uparrow i}(m)$ is the stiffness-dependent rate of nuclear import or protein synthesis for component $i$, and $k_{\downarrow i}(m)$ is the stiffness-dependent rate of nuclear export or protein degradation for component $i$. $\overrightarrow{x^{ref}}$ with elements $x_{i=1..n}^{ref}$ is a vector of arbitrary reference concentrations such that the steady-state concentration $x_i = x_i^{ref}$ when stiffness $m = m_0$. $c_{ij}$ are elements of the cooperativity matrix $C$ which we define to be the matrix of activity coefficients which describe the degree of cooperation or anti-cooperation between different elements of $\vec{x}$. Additional cooperativity matrices corresponding to more complex interactions between elements of $\vec{x}$ can be defined and added to (s1). This defines a coupled set of rate equations for each mechanosensitive phenotype marker of the cell which has a unique steady state depending on the value of $m$.

Next, we consider contributions from positive feedback loops to the dynamics of each element of $\vec{x}$. Positive feedback loops arise from active transcription which assists phenotypic shifts that promote further transcription. We add a Hill relation with coefficient $\beta$ to each equation for $\dot{x}_i$

$$\dot{x}_i = k_{\uparrow i}(m)\left(x_i^{ref} - x_i\right) - k_{\downarrow i}(m)x_i + \sum_j c_{ij}(m)x_i x_j + \alpha_i(\overrightarrow{y_k}, z)\frac{x_i^{\beta}}{x_i^{\beta} + 1} \quad \text{(s2)}$$

scaled by sensitivity $\alpha_i(\overrightarrow{y_k}, z)$, which are components of the sensitivity vector $\vec{\alpha}$. $\overrightarrow{y_k}$ is a vector of concentrations of global transcriptional participants, which may or may not all be explicitly mechanosensitive; $\overrightarrow{y_k}$ contains all the components of $\vec{x}$, and therefore has dependence on ECM stiffness $m$. $z$ is a label of the chromatin conformational state, which can be thought of as the single-cell Hi-C map of chromatin contacts; $z$ also depends on $m$ *via* physical changes to the nucleus initiated by the LINC complex (*55*, *69*). Altogether, the chromatin state $z$ and the global transcriptional cofactors $\overrightarrow{y_k}$ determine how effectively the mechanosensitive components of $\vec{x}$ can self-reinforce.

Section II: Derivation of Nonlinearly Dynamic Reinforcement Sensitivity

Each element $\alpha_i(\overrightarrow{y_k}, z)$ can be written as a sum expansion of reinforcement matrices $A^{(n)}$ multiplying $\overrightarrow{y_k}$ and $z$:

$$\alpha_i = \sum_k a_{ik}^{(1)} y_k + a_{iz}^{(1)} z + \sum_k a_{ikz}^{(2)} y_k z + \sum_{k,l} a_{ikl}^{(2)} y_k y_l + \sum_{k,l} a_{iklz}^{(3)} y_k y_l \, z \, ... \quad \text{(s3)}$$

where $a^n$ are elements of reinforcement matrices $A^n$ with dimension $n+1$. These matrix elements are weights which represent the degree to which each component of the global transcriptional environment or the global chromatin conformational state influences the self-reinforcing capability of mechanosensitive component $x_i$. We are interested in how this self-reinforcing capability evolves over time, and we use the chain rule to write the time derivative of $\alpha_i$ as

$$\frac{d\alpha_i}{dt} = \sum\nolimits_k \frac{\partial \alpha_i}{\partial y_k}\frac{\partial y_k}{dt} + \frac{\partial \alpha_i}{\partial z}\frac{\partial z}{dt} \quad \text{(s4)}$$

Plugging s3 into s4, we arrive at

$$\frac{d\alpha_i}{dt} = \sum\nolimits_{k,l\ldots} (a_{ik}^{(1)} + a_{ikz}^{(2)} z + a_{ikl}^{(2)} y_l + a_{iklz}^{(3)} y_l z + \ldots) \frac{\partial y_k}{dt} + (a_{iz}^{(1)} + a_{ikz}^{(2)} y_k + \ldots) \frac{\partial z}{dt} \quad \text{(s5)}$$

Here, we see that the dynamics of self-reinforcement sensitivity depend on dynamics of the transcription regulatory environment and the chromatin conformation, weighted by the matrix elements of the reinforcement matrices $A_i^{(n)}$. $\frac{dy_k}{dt}$ and $\frac{dz}{dt}$ are equivalent to timescales $\tau$ for each transcriptionally active component and the chromatin conformation, respectively, and generally can depend on $x_i$ and $m$. The coefficients $a_{ik}^{n}$ are generally non-linear functions of $y_k$, analogously for $a_{zk}^{(n)}$ depending on $z$. Given sufficient data to populate the partial derivative relations and reinforcement matrices in equation s5, the steady-state dynamics of cellular plasticity can be completely specified through this framework. However, this relies on highly detailed, time-dependent mechanistic knowledge which is far beyond the scope of current experimental or simulation techniques. Rather than estimate all these individual relationships with placeholder coefficients or linear rate equations, we separate the components of s5 into two timescales and perform an averaging to distill out complexity while preserving phenomenological features. Since the vector $y_k$ contains transcriptionally active components of $x$ and therefore depends on the mechanical priming program $m(t)$, we know that some terms in s5 will change on the same timescale as $x_i$ and that this timescale is an upper bound for $\frac{d\alpha_i}{dt}$. We make an arbitrary but phenomenologically justified choice of $\vec{c_i}\frac{m^\zeta}{m^\zeta+1}$ to represent these fast non-linear processes, where the time dependence originates from $m(t)$, and gather the slower terms into a separate term $\alpha_i(t_{slow})$. This term still retains $x_i$ dependence and $m$ dependence from components of $\frac{dy_k}{dt}$ and $\frac{dz}{dt}$ but contains all the slower processes in these vectors (introduced as $\tau_s \frac{m}{m_0}$ and $\tau_f$) as well as the nonlinear scaling originating from the coefficients of $A_i^{(n)}$ (introduced as $\alpha \exp\,(-\frac{x}{x_{ref}})$). Splitting $\alpha_i(t_{slow})$ into a piecewise function by region is a phenomenological choice but reflects the fact that different terms favoring an increase, decrease, or equilibration of the sensitivity will dominate depending on the magnitude of the external mechanical signal. Finally, when we perform an averaging over the components $x_i$ in the main text, the system of equations described in s5 collapse into a single equation below with two terms in each region describing both fast and slow dynamics of mechanosensitive self-reinforcement.

$$\frac{d\alpha}{dt} = \begin{cases} -\dfrac{\alpha-\alpha_0}{\tau_f} + c\dfrac{m^\zeta}{m^\zeta+1}, & I \\ \dfrac{\alpha}{\tau_s}\dfrac{m}{m_0}\exp -\dfrac{x}{x_{ref}} + c\dfrac{m^\zeta}{m^\zeta+1}, & II \\ -\dfrac{\alpha}{\tau_s}\dfrac{m}{m_0}\exp -\dfrac{x}{x_{ref}} + c\dfrac{m^\zeta}{m^\zeta+1}, & III \end{cases} \quad \text{(s6)}$$

<u>Section III: Linear Dynamics used for Noise Simulations</u>

Linear Dynamics for $\frac{d\alpha}{dt_{slow}}$ (Figure S5):

$$\frac{d\alpha}{dt_{slow}} = \begin{cases} -\dfrac{\alpha - \alpha_0}{\tau_f}, & I \\ \dfrac{\alpha}{\tau_s}, & II \\ -\dfrac{\alpha}{\tau_s}, & III \end{cases}$$

| **Reference** | **Cell Type** | **Priming Stiffness (kPa)** | **Priming Time (days)** | **Memory Stiffness (kPa)** | **Memory Time (days)** | **Memory marker** |
|---|---|---|---|---|---|---|
| Yang et al. (*6*) | hMSC | 10 | 1, 7, 10 | 2 | 1, 5+, 10+ | YAP, RUNX2 |
| Balestrini et al. (*4*) | Fibroblasts | 25, 100 | 14 | 5 | 14+ | α-SMA |
| Xi et al. (*5*) | rMSC | 100 | 21 | 5 | 14+ | α-SMA, miR-21 |
| Nasrollahi et al. (*7*) | mcf10a<br>A431<br>Mcf7 | 50 | 3 | 0.5 | 3+ | pMLC, YAP, migration speed |
| Watson et al. (*66*) | SUM159 | 8 | 7 | 0.5 | 2 to 7 | RUNX2, migration speed |
| Dunham et al. (*10*) | ASC | 5 | 14 | 100 | 7 to 14 | Cell area, α-SMA |

**Table S1.** Summary of experimental data collected on mechanical memory.

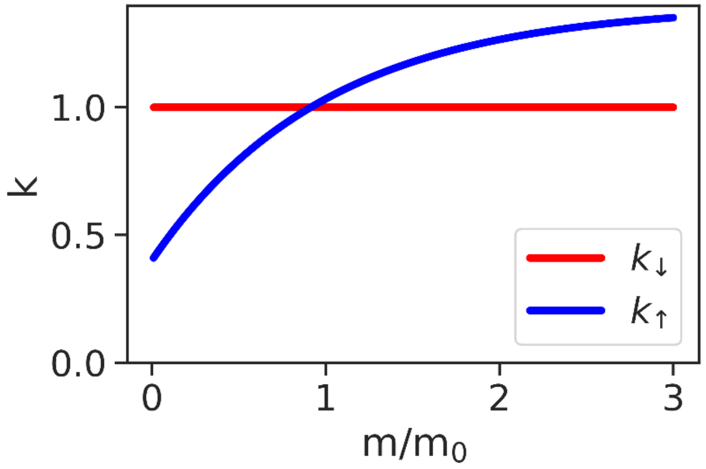


**Fig. S1.**

Mechanosensitivity of synthesis and nuclear localization of stiff-correlated transcription factors (blue) and countering degradation and nuclear export (red).

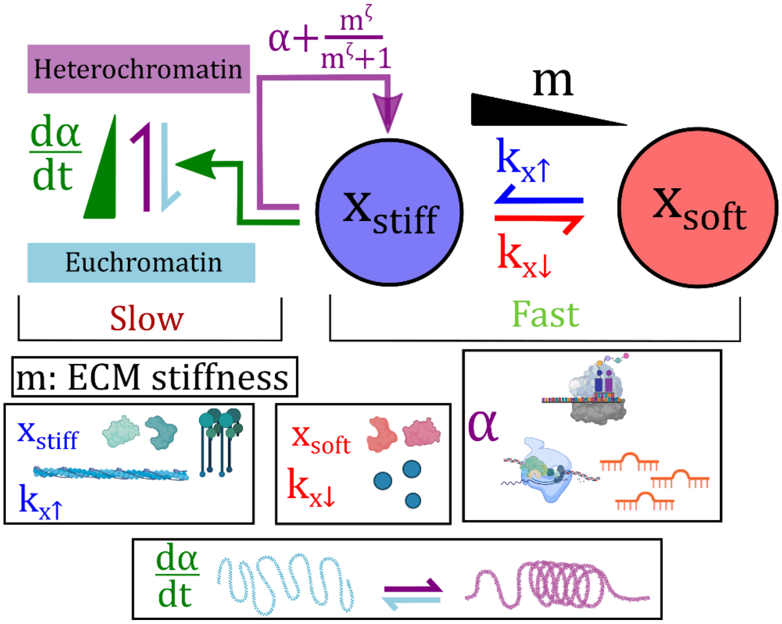


**Fig. S2.**

**Circuit diagram of the dynamic mechanical memory model.** x represents the mechanoactivated phenotype of the cell including nuclear localized transcription factors such as YAP, RUNX2, and MKL-1. m represents ECM stiffness. Nuclear x self-reinforces with transcriptional efficiency $\alpha + \frac{m^{\zeta}}{m^{\zeta}+1}$; the first term represents signal-driven effects on transcription, while the second term represents LINC-driven physical processes. $\frac{d\alpha}{dt}$ gives the change of the efficiency of this self-reinforcement over time *via* a modified transcriptional landscape.

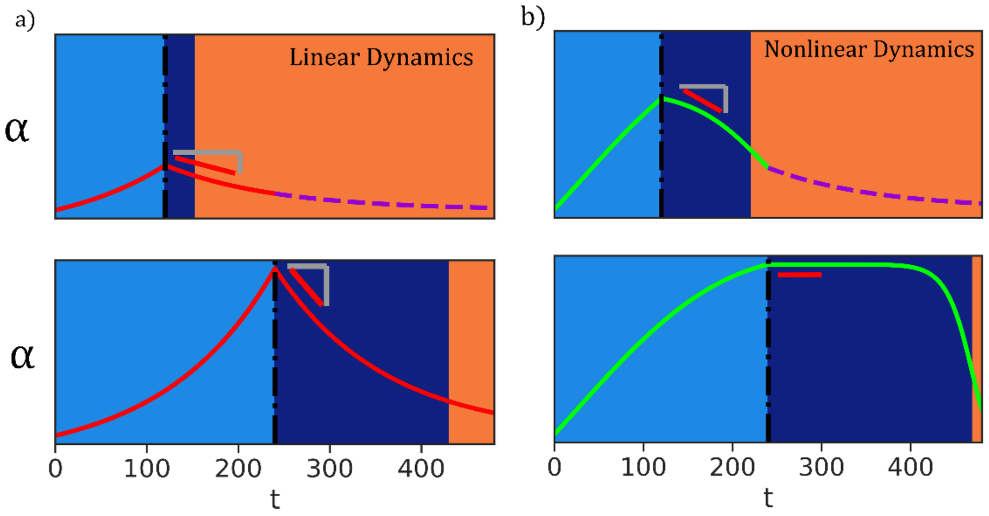


**Fig. S3.**

Comparison of linear and non-linear dynamics of $\alpha$. **(A)** Linear dynamics. **(B)** Nonlinear dynamics. While shorter primes (top row) lead to less memory (width of dark blue region) than longer primes (bottom row) for both (A) and (B), the initial rate of memory dissipation is much faster in (A) than in (B); (B) matches better with experiment.

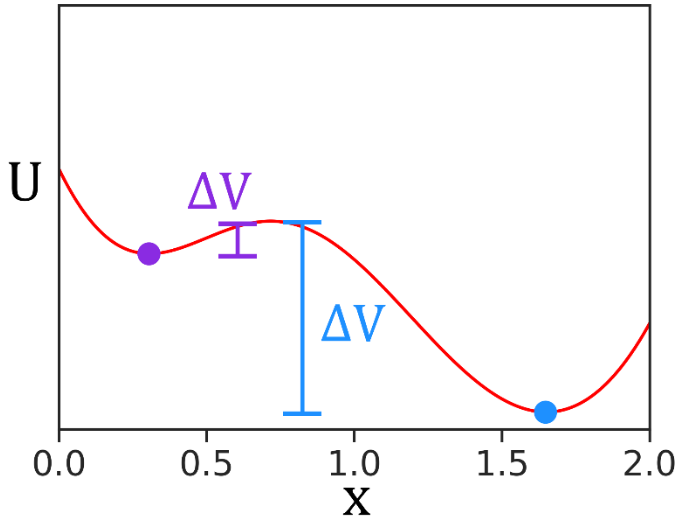


**Fig. S4.**

Illustration of local minima in region III of the phase diagram shown in Figure 2. If random fluctuations perturb the state from the steady state minimum, the population in the shallow well (purple) will be reduced relative to the population in the deeper well (blue).

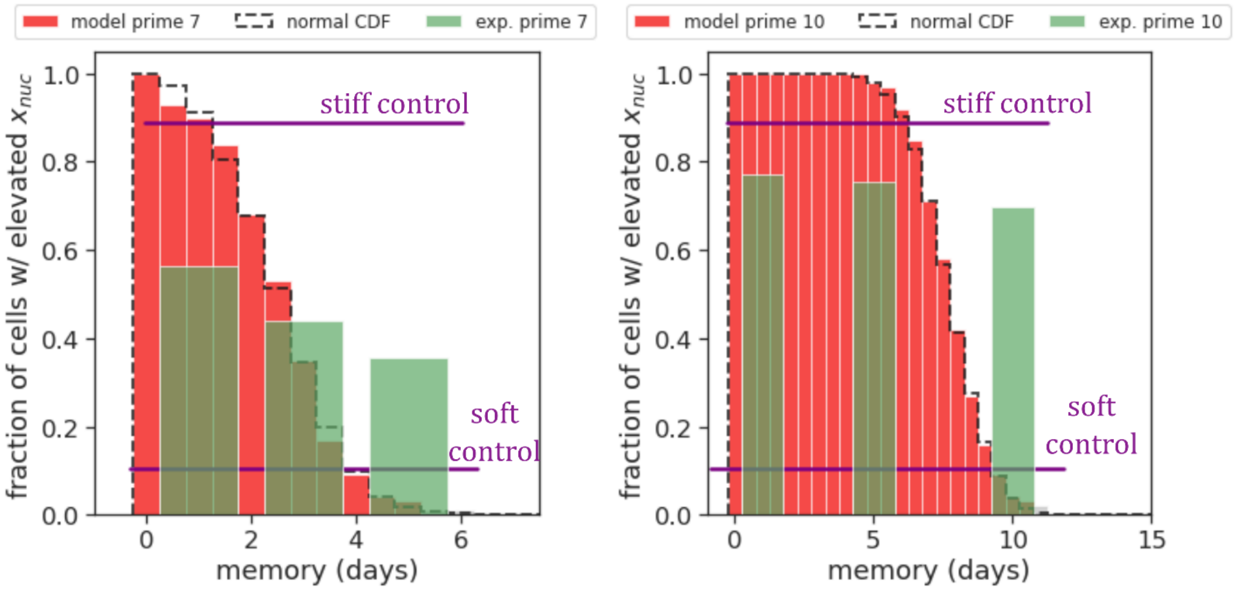


**Fig. S5.**

Noise study with linear dynamics, where dependence of the slow varying component of $\frac{d\alpha}{dt}$ on $x$ is removed. Gaussian noise applied in this situation leads to a normal distribution of memory time, in contrast with the non-normal distribution of nonlinear dynamics and in contrast with experimental results (green bars).

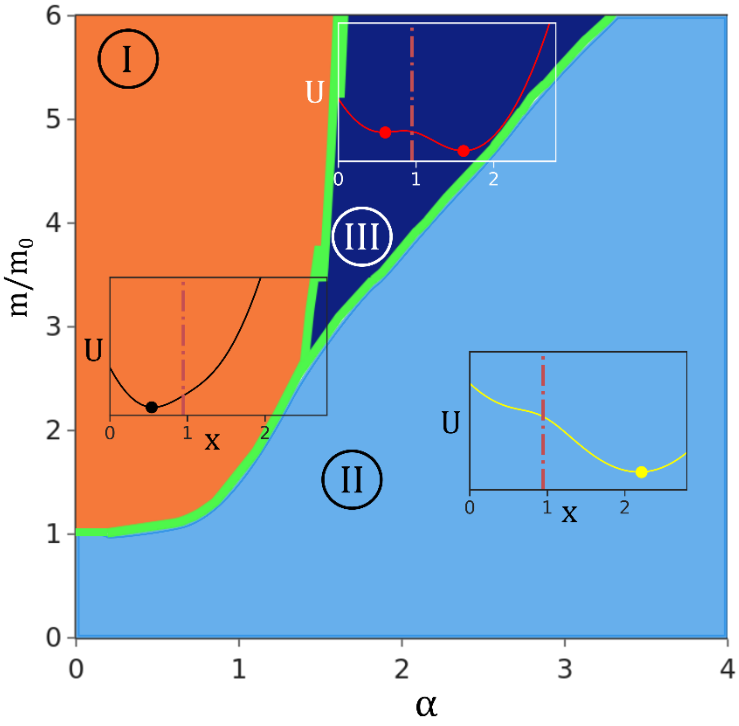


**Fig. S6.**

Analogous phase diagram of the model for soft-activated genes. In the model, the mechanoactivation profile / mechanical signaling is reversed by flipping $\frac{m}{m_0}$ to $\frac{m_0}{m}$, so that $\frac{dx}{dt}$ increases when stiffness is reduce. In this case, high $x$ corresponds to activity of soft-correlated phenotypic genes and transcription factors. $\alpha$ now represents positive reinforcement for gene expression correlating with a soft phenotype.
